# Comparative transcriptomics of iPSC-derived cardiomyocytes in a high and a low altitude population

**DOI:** 10.1101/2025.06.04.657917

**Authors:** OA Gray, DB Witonsky, A Di Rienzo

## Abstract

Tibetan adaptation to high-altitude hypoxia remains a classic example of Darwinian selection in humans. To identify adaptive traits that might have evolved in Tibetans in response to long-term exposure to hypoxia, we previously established a library of induced pluripotent stem cells (iPSCs), derived from Tibetan and Han Chinese individuals, as a robust model system for the exploration of condition-specific molecular and cellular responses. We used this system to characterize and compare the transcriptome of iPSC-derived endothelial cells and found that angiogenesis, energy metabolism and immune pathways differ between the cell lines from these two populations. Here, we harness the same experimental system to characterize and compare the transcriptome of iPSC-derived cardiomyocytes in Tibetan and Han Chinese in hypoxia. We find that several pathways, such as the hypoxia, myogenesis and glycolysis pathways, are significantly enriched for differentially expressed genes across populations. These pathways are candidate targets of natural selection due to exposure to the high-altitude hypoxic environment and point to adaptive cardiac traits such as sustained cardiac function in hypoxia. A better understanding of these adaptations may offer insights into novel therapeutic strategies for hypoxia-related cardiovascular conditions, such as pulmonary hypertension and ischemic heart disease.

## Introduction

Tibetan populations provide a well-characterized example of Darwinian selection in humans, having been exposed to high altitude hypoxia for over 10,000 years^1–3^. Over time, this selective pressure has resulted in a suite of adaptative phenotypes that are not shared between Tibetans and closely-related acclimatized lowlanders, such as the Han Chinese, and/or other long-term high-altitude residents, such as Andeans^4–7^. Most notably, compared to East Asian lowlanders at high altitude, Tibetans exhibit lower rates of pregnancy complications (e.g. preeclampsia) and higher birth weights, a blunted erythropoietic response to hypoxia, no or modest pulmonary hypoxic vasoconstriction, and low rates of hypoxic pulmonary hypertension (PH)^7–10^. They also have lower rates of chronic mountain sickness (CMS) compared to Andean highlanders ^6,7,11^.

Selection scans in populations of Tibetan ancestry have identified several signals of strong selection at individual loci, among which the most well replicated ones are at the genes *EPAS1* and *EGLN1*^12–14^. Both genes code for key regulators of the transcriptional response to hypoxia. The selected alleles at *EGLN1* are nonsynonymous though it is still debated whether they are gain- or loss-of-function variants^15,16^. In contrast, all alleles on the selected haplotype at *EPAS1* are noncoding and disrupt multiple enhancers resulting in a dampened transcriptional response to hypoxia across multiple cell and tissue types^17–19^. These alleles have been associated with reduced hemoglobin concentration^8,9,12,20^ and with low pulmonary vasoconstriction^17^, both measured at high altitude. Polygenic adaptation tests also revealed signals for reproductive and cardiovascular traits, including better reproductive outcomes^8,10^, higher pulse^8^, slower blood clotting^10^, lower serum albumin and triglyceride levels^10,21^, greater right ventricular inner diameter^10^, higher lactate dehydrogenase and eosinophil count^21^.

Most adaptive traits have been identified through field work by measuring organismal phenotypes at high altitude in both Tibetans and acclimatized lowlanders. An alternative approach compares molecular traits, e.g. transcript levels, in cohorts of Tibetans and of a closely related low-altitude population, e.g. Han Chinese, both measured in hypoxic conditions. This approach requires cell culture models for the populations and the tissues of interest. The collection of primary tissue samples presents a number of logistical challenges and, for this reason, has been limited to easily accessible cell types, such as umbilical vein endothelial cells (HUVECs)^17^, peripheral blood cells^20^, or lymphoblastoid cell lines (LCLs)^22^. While valuable resources, these cell types alone cannot address the impacts of adaptation across diverse human tissues and physiological systems. To circumvent these limitations, we recently established a panel of induced pluripotent stem cells (iPSC) from individuals of Tibetan or Han Chinese ancestry^21^, which can be differentiated into cell types of interest that are not easily accessible *in situ*. Because the vascular endothelium is a primary mediator of the response to hypoxia^23,24^ and has been directly implicated in Tibetan adaptation^17–19^, we previously used this panel to identify differentially expressed (DE) genes in iPSC-derived endothelial cells between Tibetans and Han Chinese. This work pointed to candidate adaptive phenotypes in Tibetans that involve metabolic and inflammatory processes.

Here, we use the same panel to extend our investigation of transcriptional adaptations to hypoxia in iPSC-derived cardiomyocytes. The heart plays a critical role in oxygen delivery and utilization, and molecular adaptations in cardiomyocytes are likely to reflect key mechanisms in Tibetan high-altitude physiology. In turn, elucidating these cardiac adaptations may shed new light on hypoxia-related disorders such as pulmonary hypertension or heart failure. We find several strongly DE genes in cardiomyocytes, such as *SLC6A4* and *COX6A2*, with links to adaptive cardiac traits such as PH and myocardium remodeling, respectively. Additionally, we find that the Hypoxia, Myogenesis and Glycolysis pathways are enriched for DE genes across populations, hence, may have been shaped by the selective pressures of the high-altitude environment.

## Results and Discussion

### iPSC panel & differentiation

We previously established a panel of 20 sex-matched iPSC lines from individuals of Tibetan and Han Chinese ancestry by reprogramming their LCLs^25^. The panel was specifically designed to identify candidate adaptive molecular traits that, on average, differ between Tibetans and a closely related population without a history of long-standing residence at high altitude. While the Tibetan cohort was sampled in the Chicago area, henceforth referred to as the Tibetan Alliance of Chicago (TAC) samples, the Han Chinese (CHB) LCLs were obtained from the 1000 Genome (1KG) project^22^. Importantly, to avoid technical confounders, the 20 LCLs were reprogrammed into iPSCs in batches, balanced by population and sex, and subjected to multiple quality control (QC) steps, as described in Gray et al., 2025^21^.

These 20 iPSC lines were differentiated into cardiomyocytes -- using the protocol described in Knowles et al., 2018^26^ -- in 7 batches containing either three or four cell lines. Each batch included approximately equal numbers of TAC and CHB lines, as well as an equal distribution of males and females. Each batch of iPSC-CM lines was cultured in hypoxic conditions (1% O_2_) for 48 hrs, then checked for viability (see Table S1). Samples within each batch were pooled prior to single-cell isolation and RNA sequencing (scRNA-seq; see Methods). Individual lines within pooled samples were deconvoluted using the genotype data and subjected to additional QC steps, including doublet cell removal, filtering by number of expressed genes (1000 < feature counts < 5000) and mtDNA counts (<10%) resulting in 11,632 and 11,623 high quality cells in total from the CHB and TAC lines, respectively, which corresponded to 54% of the cells prior to QC. Data were subsequently normalized, differences in cell cycle phase among cells were regressed out, and batch-specific effects were corrected for (see Methods). Cells were then clustered based on their transcriptional profiles leading to the identification of 11 distinct clusters (Fig. 1A). To identify the cells with cardiac features, we looked at the expression levels of known cardiac marker genes^27^ and found that clusters 1, 3 and 4 consistently express cardiac markers (Fig. 1B); we will refer to these clusters as CM clusters. We also used the top 400 DE genes across clusters to perform gene set enrichment analysis with the Human Protein Atlas Normal Tissue database^28^. This analysis showed that the same three clusters were highly enriched for genes expressed in cardiomyocytes.

**Fig 1.**
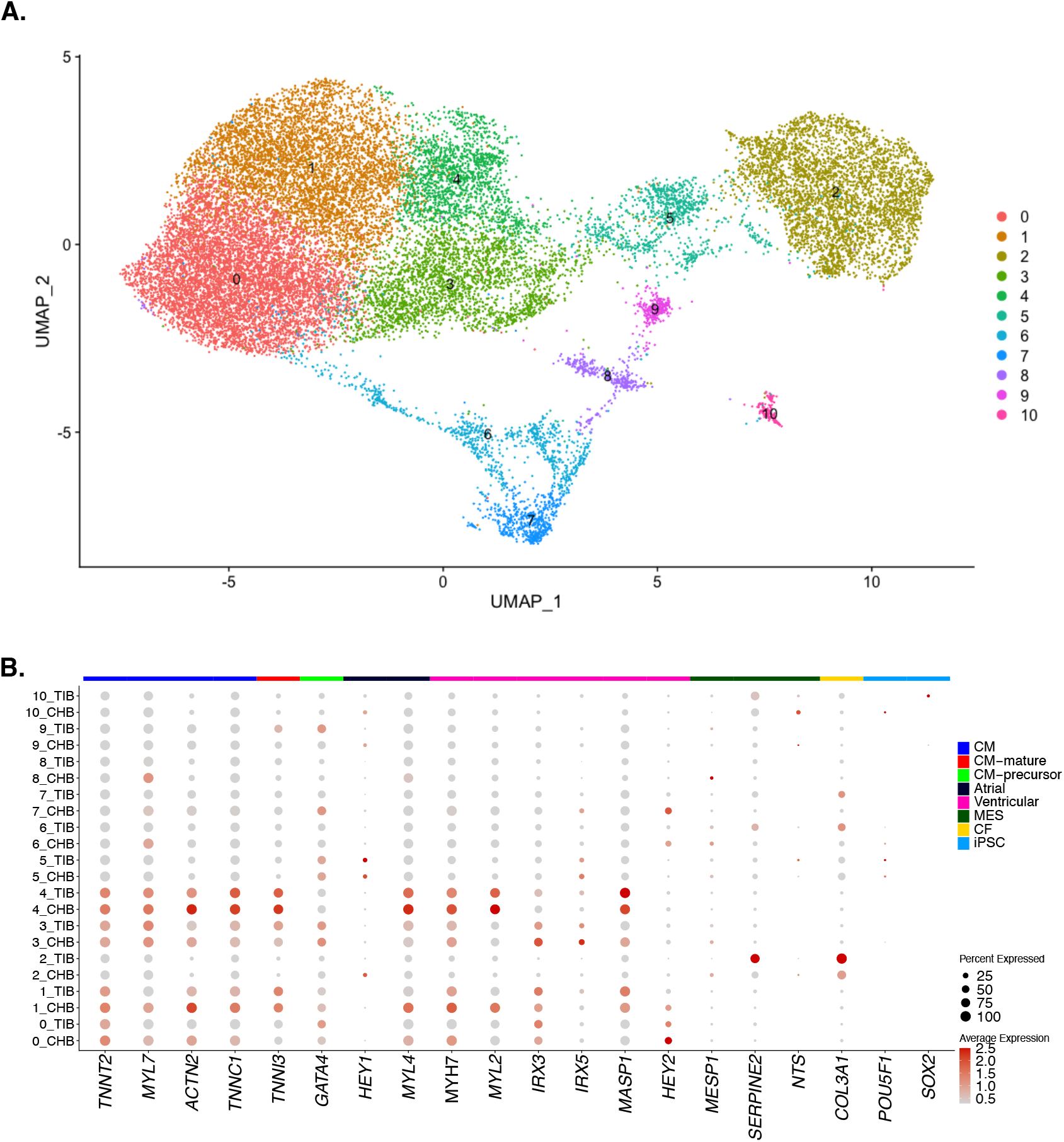
A) UMAP plot of iPSC-CMs colored by cluster after Harmony reduction. B) Expression of cardiac marker genes across the 11 cell clusters (labeled 0–10) identified in the scRNAseq data shown in A. Each row of the dot plot shows the cluster number and the population of origin, whereas the size and color of the dots indicate, respectively, the percent of cells expressing that gene and the average expression level. Across both populations, clusters 1, 3 and 4 robustly express cardiomyocyte marker genes.

### Detection of Transcriptional Differences Between Low- and High-Altitude Populations

Next, we tested for differences in transcript levels between CHB and TAC lines to identify molecular phenotypes that evolved in Tibetans during adaptation to the high-altitude environment. These molecular traits may offer insights into how the cardiovascular system was shaped by natural selection due to hypoxia exposure and point to novel beneficial organismal traits to be tested in future field studies.

For differential expression (DE) analyses, we aimed to test the pooled cells from the three CM clusters across all individuals from each population; we will refer to this pool as the CM pool. However, we noticed that there was great inter-individual variation in cell number, with a subset of individuals contributing most of the cells assigned to a given cluster or subset of clusters (Table 1). Therefore, to minimize the possible confounding of DE with individual cell count heterogeneity, we pruned our data set to the 10 individuals, 5 CHB and 5 TAC, contributing the largest number of cells to the CM pool, with 408 cells being the lowest (Table 2).

**Table 1.** Number of cells per cluster in each individual cell line and pooled by population.

| Cell line ID | Cluster |  |  |  |  |  |  |  |  |  |  |
| --- | --- | --- | --- | --- | --- | --- | --- | --- | --- | --- | --- |
|  | 0 | 1 | 2 | 3 | 4 | 5 | 6 | 7 | 8 | 9 | 10 |
| NA18528 | 133 | 138 | 37 | 51 | 19 | 22 | 6 | 5 | 1 | 9 | - |
| NA18531 | 60 | 194 | 27 | 38 | 118 | 43 | 10 | 9 | 14 | 4 | 1 |
| NA18557 | 520 | 633 | 39 | 12 | 129 | 9 | 59 | 91 | 12 | 17 | 6 |
| NA18596 | 19 | 15 | 729 | 325 | 157 | 94 | 89 | 24 | 52 | 38 | 10 |
| NA18606 | 1,032 | 613 | 218 | 402 | 165 | 165 | 132 | 35 | 53 | 19 | 3 |
| NA18608 | 1 | - | 69 | 1 | 3 | 6 | 9 | 2 | 4 | - | - |
| NA18614 | 1,104 | 429 | 17 | 120 | 94 | 6 | 122 | 123 | 5 | 2 | 38 |
| NA18619 | 29 | 22 | 358 | 556 | 193 | 42 | 38 | 21 | 11 | 100 | - |
| NA18633 | 58 | 93 | 17 | 32 | 6 | 13 | 5 | - | 1 | 6 | - |
| NA18748 | 10 | 51 | 46 | 261 | 164 | 216 | 90 | 10 | 122 | 20 | 31 |
| TAC252 | 1,654 | 1,226 | 58 | 158 | 274 | 55 | 70 | 47 | 19 | 2 | 28 |
| TAC462 | 232 | 249 | 1 | 28 | 56 | 46 | 14 | 15 | 9 | 3 | - |
| TAC540 | 334 | 57 | 42 | 44 | 17 | 12 | 28 | 16 | 5 | 2 | - |
| TAC569 | 33 | 6 | 144 | 361 | 48 | 87 | 30 | 12 | 22 | 17 | 29 |
| TAC750 | 23 | 32 | 39 | 10 | 6 | 13 | 3 | 2 | 3 | - | - |
| TAC801 | 989 | 395 | 2 | 4 | 9 | 2 | 18 | 47 | 2 | 1 | - |
| TAC841 | 40 | 5 | 18 | 165 | 33 | 37 | 10 | - | 2 | 4 | 2 |
| TAC859 | 36 | 539 | 21 | 14 | 44 | 36 | 17 | 26 | 11 | 9 | 1 |
| TAC86 | 18 | 2 | 2,027 | 48 | 7 | 1 | 143 | 90 | 16 | 4 | 10 |
| TAC988 | 387 | 390 | 18 | 22 | 131 | 28 | 13 | - | 7 | - | 1 |
| CHB pool | 2,966 | 2,188 | 1,557 | 1,798 | 1,048 | 616 | 560 | 320 | 275 | 215 | 89 |
| TAC pool | 3,746 | 2,901 | 2,370 | 854 | 625 | 317 | 346 | 255 | 96 | 42 | 71 |

**Table 2.**
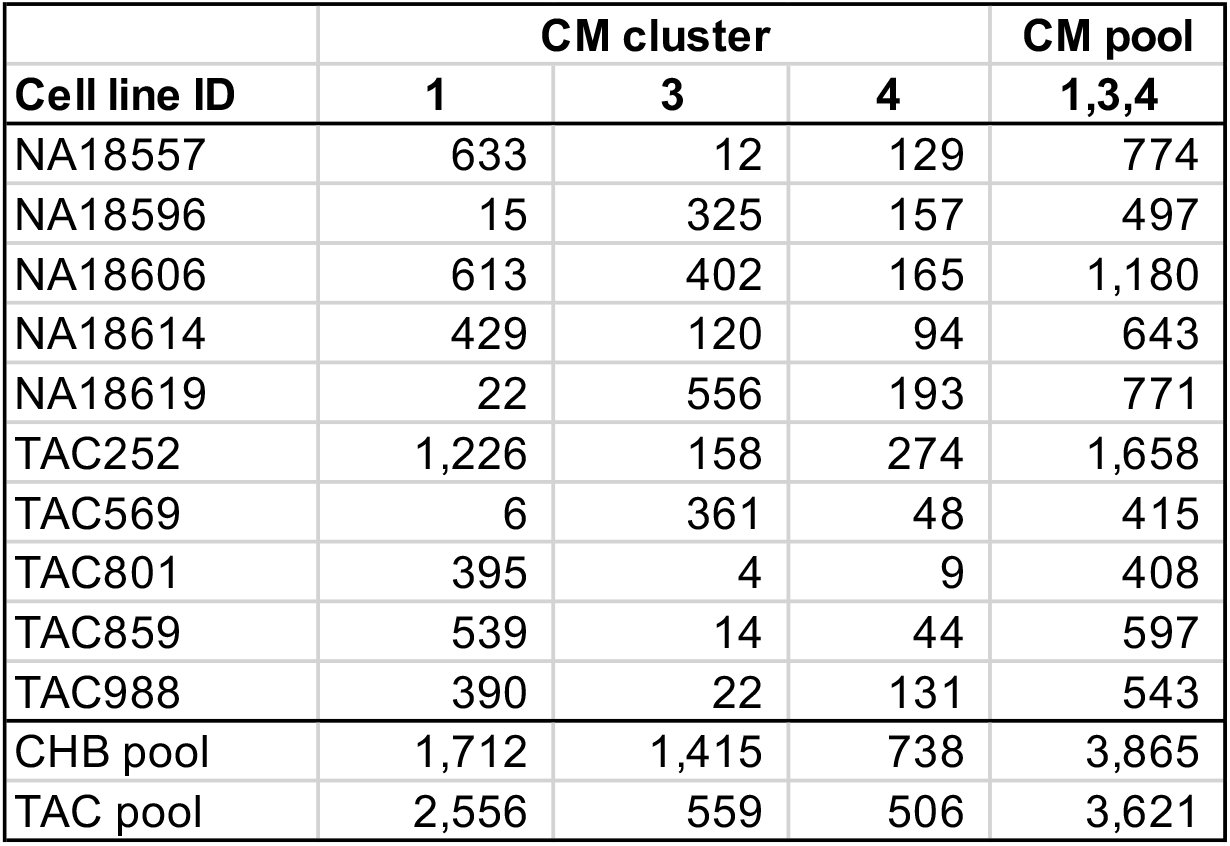
Number of cells in each CM cluster and in the CM pool (clusters 1, 3, 4) in the 10 lines used for DE analysis and in the two population pools.

| **Cell line ID** | **CM cluster** |  |  | **CM pool** |
| --- | --- | --- | --- | --- |
|  | **1** | **3** | **4** | **1,3,4** |
| NA18557 | 633 | 12 | 129 | 774 |
| NA18596 | 15 | 325 | 157 | 497 |
| NA18606 | 613 | 402 | 165 | 1,180 |
| NA18614 | 429 | 120 | 94 | 643 |
| NA18619 | 22 | 556 | 193 | 771 |
| TAC252 | 1,226 | 158 | 274 | 1,658 |
| TAC569 | 6 | 361 | 48 | 415 |
| TAC801 | 395 | 4 | 9 | 408 |
| TAC859 | 539 | 14 | 44 | 597 |
| TAC988 | 390 | 22 | 131 | 543 |
| CHB pool | 1,712 | 1,415 | 738 | 3,865 |
| TAC pool | 2,556 | 559 | 506 | 3,621 |

DE analysis between CHB and TAC lines was performed in a pseudo-bulk manner by first aggregating read counts of the CM clusters in the single cell analysis for each sample. We tested genes for DE across populations with the design model ∼population+batch using DEseq2^29^. DE analysis across TAC and CHB lines was then performed using all cells in the CM pool with varying cell numbers across individuals as well as a subsample of only 408 cells for each individual to assess the robustness of our results to varying cell numbers. As shown in Fig. 2, we found a strong correlation between the *Z*-scores from the DE analysis including all cells vs. the 408-cell subsample (r^2^=0.985). Therefore, we decided to use the results using all cells in subsequent analyses.

**Fig 2.**
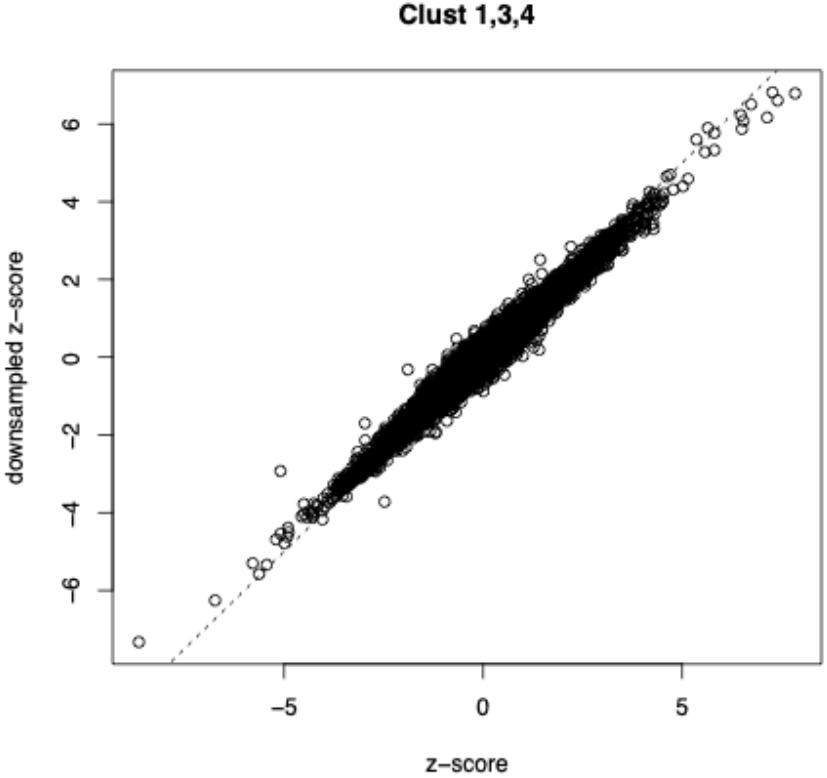
Correlation between the results of the DEseq2 analysis in all cells and a sample of 408 cells for each individual in the CM pool. The horizontal axis shows the *Z*-scores of the DE test in all cells while the vertical axis shows the *Z*-scores for the subsample of 408 cells per line.

We found 9 genes that are shared among the top 10 most significantly DE across both analyses (listed in Table 3), i.e. all cells and 408 cells (Tables S2-3). A negative log_2_ fold change in transcript levels (LFC) indicates that the gene is expressed at lower levels in TAC compared to CHB lines, *vice versa* for a positive LFC. Among these DE genes, *SLC6A4*, which codes for the serotonin transporter, is one of the most interesting. While serotonin is often associated with the nervous system, it also plays a crucial role in the cardiovascular system affecting various aspects of cardiac function, such as heart rate, contractility, and vascular tone^30^. The serotonin transporter helps regulate the availability of serotonin by facilitating its reuptake, thereby modulating the local concentrations of serotonin^31^. In mice, *Slc6a4* is expressed in embryonic, but not adult myocardium, and in cardiac valves; moreover, *Slc6a4* knock out mice show marked cardiac phenotypes, such as cardiac valvulopathy and myocardial fibrosis^32^. Additionally, there is clear evidence linking the serotonin transporter with hypoxia-induced PH^33^. Mice lacking the transporter are relatively resistant to hypoxia-induced PH and those overexpressing it demonstrate a pathology that resembles pulmonary hypoxia, namely pulmonary arterial smooth muscle cell hypertrophy, and cardiac hypertrophy^34,35^. In addition, the selective inhibitors of the serotonin transporter, such as fluoxetine, have been shown to protect against hypoxic PH^36^, a relatively common condition in acclimatized lowlanders at high altitude, which is rare in Tibetans. Consistent with the idea that low levels of *SLC6A4* expression might be advantageous at high altitude, we find that this gene is strongly and significantly down-regulated in Tibetans (Table 3).

**Table 3.**
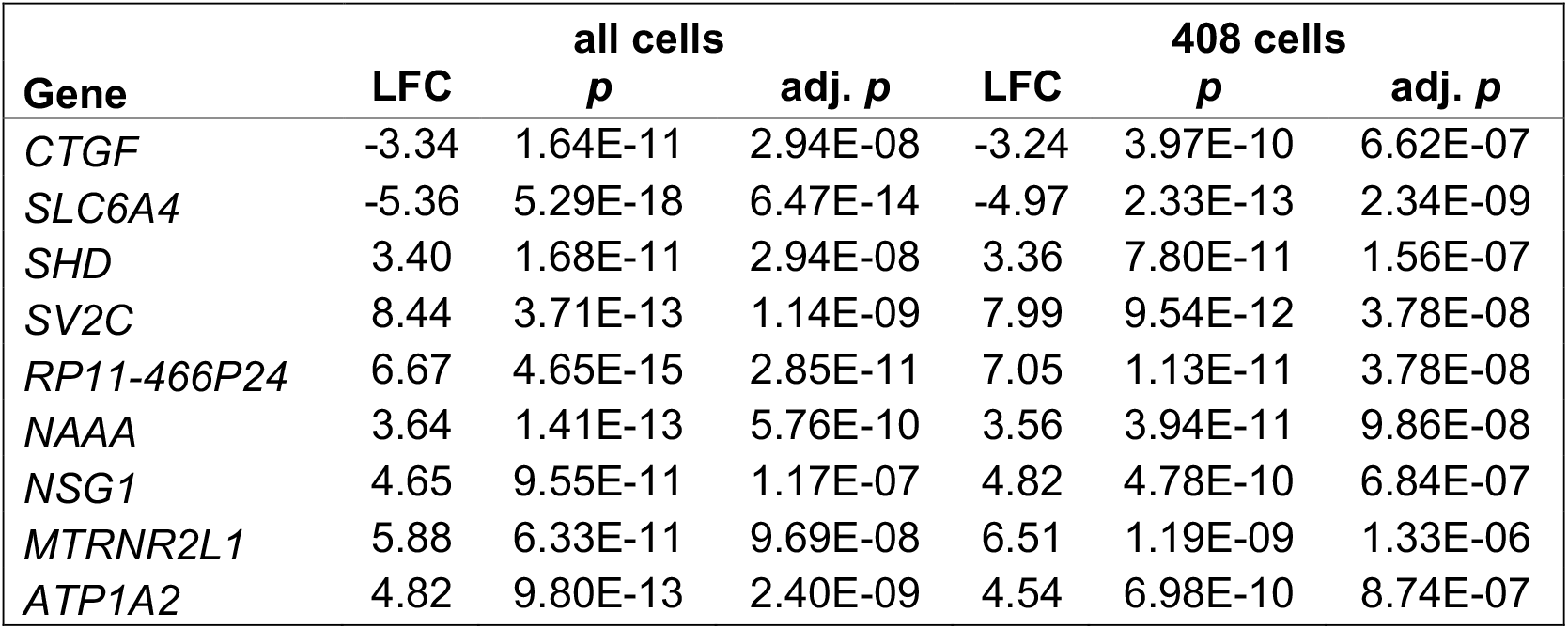
Shared top DE genes between populations in the DEseq2 analysis including all cells or only 408 cells for each individual.

| **Gene** | **LFC** | **all cells** |  | **LFC** | **408 cells** |  |
| --- | --- | --- | --- | --- | --- | --- |
|  |  | ***p*** | **adj. *p*** |  | ***p*** | **adj. *p*** |
| *CTGF* | -3.34 | 1.64E-11 | 2.94E-08 | -3.24 | 3.97E-10 | 6.62E-07 |
| *SLC6A4* | -5.36 | 5.29E-18 | 6.47E-14 | -4.97 | 2.33E-13 | 2.34E-09 |
| *SHD* | 3.40 | 1.68E-11 | 2.94E-08 | 3.36 | 7.80E-11 | 1.56E-07 |
| *SV2C* | 8.44 | 3.71E-13 | 1.14E-09 | 7.99 | 9.54E-12 | 3.78E-08 |
| *RP11-466P24* | 6.67 | 4.65E-15 | 2.85E-11 | 7.05 | 1.13E-11 | 3.78E-08 |
| *NAAA* | 3.64 | 1.41E-13 | 5.76E-10 | 3.56 | 3.94E-11 | 9.86E-08 |
| *NSG1* | 4.65 | 9.55E-11 | 1.17E-07 | 4.82 | 4.78E-10 | 6.84E-07 |
| *MTRNR2L1* | 5.88 | 6.33E-11 | 9.69E-08 | 6.51 | 1.19E-09 | 1.33E-06 |
| *ATP1A2* | 4.82 | 9.80E-13 | 2.40E-09 | 4.54 | 6.98E-10 | 8.74E-07 |

To identify biological processes that might have been shaped differently by natural selection in one of the two populations, we performed gene set enrichment analysis using the EnrichR tool^37–39^ (https://maayanlab.cloud/Enrichr/). All DE genes between populations with adjusted *p* value below 0.05 were tested for enrichment relative to the background of all expressed genes. We found that the two most significantly enriched gene sets are the Myogenesis and Hypoxia terms in the MSigDB_Hallmark, suggesting that indeed a history of long-term hypoxia exposure resulted in selective pressures that shaped Tibetan cardiomyocyte function. Specific genes in these sets are shown in Table 4; most of these genes, especially those in the Hypoxia gene set, are expressed at higher level (positive LFC) in TAC than in CHB lines (Table 5).

**Table 4.** Enriched gene sets in the MSigDB for DE genes (adj *p* < 0.05) detected by DEseq2.

| **Term** | **Set size** | **$p$** | **adj. $p$** | **DE genes in set** |
| --- | --- | --- | --- | --- |
| Myogenesis | 179 | 9.60E-05 | 3.40E-03 | *ACTA1; DTNA; SCD; AGL; DMD; SORBS1; COX6A2; CLU; HBEGF* |
| Hypoxia | 171 | 3.83E-04 | 6.70E-03 | *SCARB1; DTNA; PLIN2; ALDOC; ENO2; NDRG1; TGM2; IER3* |
| Apoptosis | 141 | 3.32E-03 | 3.03E-02 | *IFITM3; TNFRSF12A; NEDD9; ENO2; CLU; IER3* |
| Interferon Gamma Response | 145 | 3.81E-03 | 3.03E-02 | *IFITM3; BST2; ARID5B; ISG15; USP18; HLA-DRB1* |
| IL-2/STAT5 Signaling | 154 | 5.10E-03 | 3.03E-02 | *IFITM3; PLIN2; NDRG1; HOPX; DHRS3; TGM2* |
| Cholesterol Homeostasis | 68 | 5.20E-03 | 3.03E-02 | *TNFRSF12A; SCD; ALDOC; CLU* |
| Interferon Alpha Response | 78 | 8.41E-03 | 4.21E-02 | *IFITM3; BST2; ISG15; USP18* |

**Table 5.** DE genes driving the gene set enrichment in Hypoxia and Myogenesis.

| **Gene** | **LFC** | ***p*** | **adj. *p*** | **Enriched set** |
| --- | --- | --- | --- | --- |
| *ENO2* | 2.34 | 2.66E-06 | 1.12E-03 | Hypoxia |
| *TGM2* | -1.87 | 1.87E-05 | 4.54E-03 | Hypoxia |
| *PLIN2* | 1.96 | 3.30E-05 | 6.50E-03 | Hypoxia |
| *IER3* | -1.59 | 2.52E-04 | 2.69E-02 | Hypoxia |
| *NDRG1* | 1.82 | 4.50E-04 | 3.96E-02 | Hypoxia |
| *SCARB1* | 1.91 | 5.53E-04 | 4.21E-02 | Hypoxia |
| *ALDOC* | 1.55 | 6.72E-04 | 4.81E-02 | Hypoxia |
| *COX6A2* | 3.83 | 8.15E-11 | 1.11E-07 | Myogenesis |
| *ACTA1* | 2.9 | 1.67E-08 | 1.46E-05 | Myogenesis |
| *SCD* | 1.72 | 5.61E-06 | 2.15E-03 | Myogenesis |
| *HBEGF* | -2.24 | 6.62E-06 | 2.32E-03 | Myogenesis |
| *CLU* | -1.98 | 9.57E-05 | 1.39E-02 | Myogenesis |
| *AGL* | 1.82 | 9.85E-05 | 1.42E-02 | Myogenesis |
| *SORBS1* | 2.27 | 2.45E-04 | 2.68E-02 | Myogenesis |
| *DMD* | -1.89 | 4.61E-04 | 4.01E-02 | Myogenesis |
| *DTNA* | 2.04 | 1.04E-04 | 1.43E-02 | Myogenesis, Hypoxia |

To exploit the similarity of DE patterns across cell types, we used multivariate adaptive shrinkage (mashr)^40^ on the expression data from the iPSC-CMs and the published data from endothelial cells derived from the same panel of iPSC lines^21^. Mashr leverages the combination of many effects (in our case gene expression differences between TAC and CHB lines) ascertained across many conditions (in our case iPSC-CMs in hypoxia and iPSC-ECs, respectively, in hypoxia and normoxia) to model effect sharing across conditions and increase the power to detect effects shared across conditions as well as condition-specific effects. Significance in mashr is calculated in each tissue using the local false sign rate (*lfsr*).

Of the 12,835 genes included in the mashr analysis, we found that 100 were DE in all 3 data sets at *lfsr* <0.1 (Table S4). As shown in Fig. 3A, we found a total of 252 DE genes (*lfsr*<0.1) in iPSC-CMs in hypoxia, similar to the corresponding number (249) in iPSC-ECs in hypoxia. Perhaps surprisingly, the total number of DE genes in iPSC-ECs in normoxia is substantially larger (394). Interestingly, the most significant DE genes in iPSC-CMs were not significant in iPSC-ECs in either normoxia or hypoxia; these genes included the top DE genes detected by DEseq2 in iPSC-CMs (i.e. *SLC6A4, SV2C, NAAA, CTGF, MTRNR2L1, UNC5B-AS1, CTNND2*). This finding suggests that a subset of cardiovascular adaptations to hypoxia are indeed specific to the heart.

**Fig. 3.**
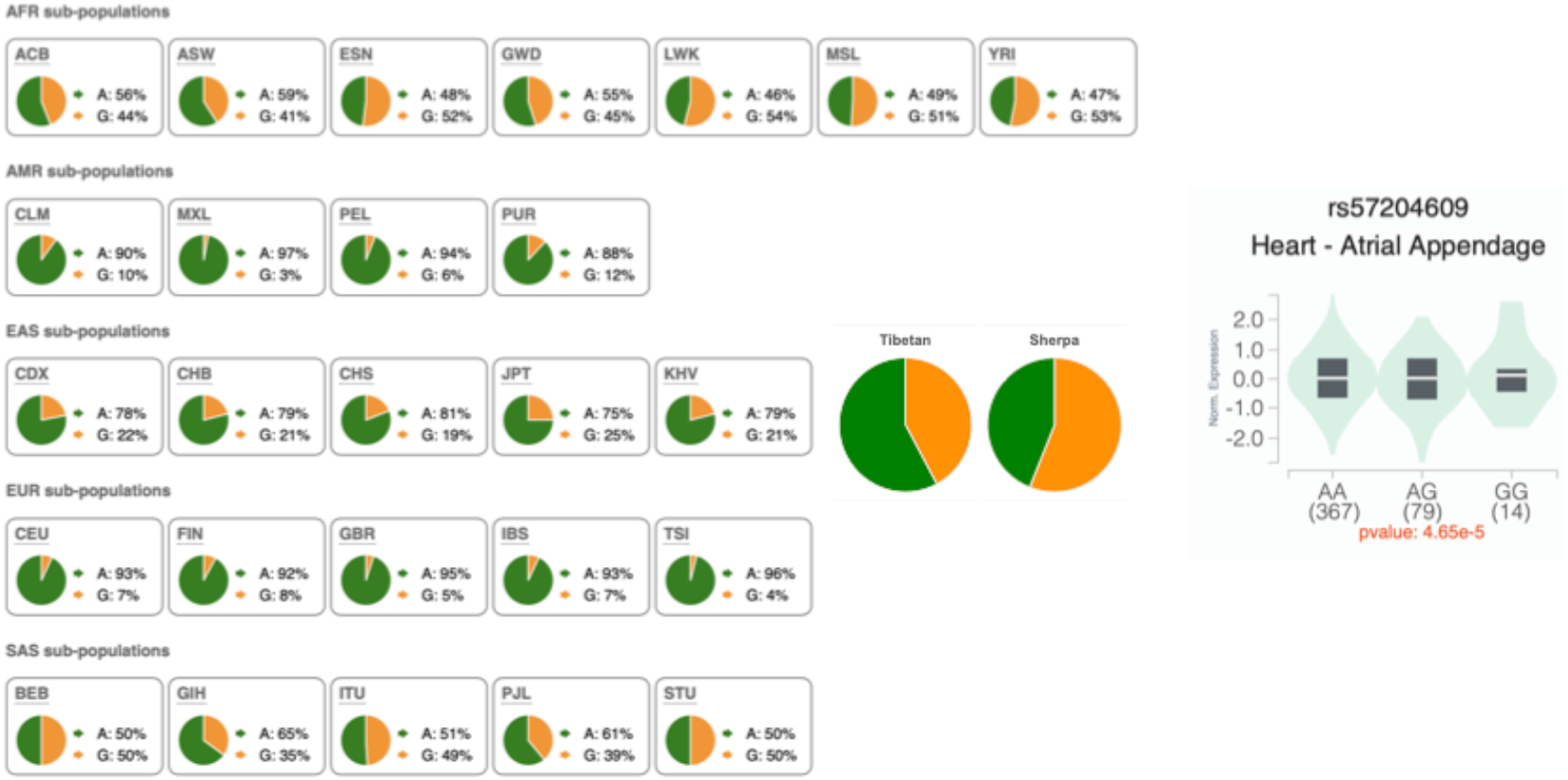
A) Allele frequencies of SNP rs57204609 in the 1000 Genomes Project Phase 3 and in Tibetan and Sherpa samples from Jeong et al. (2018)^8^. B) Violin plot for rs57204609 genotypes and *COX6A2* transcript levels in the Heart Atrial Appendage from the GTEx Analysis Release V10.

We performed gene set enrichment analysis using the DE genes detected by mashr (see Tables S5-7 and Fig. 3B). By including all the DE genes (*lfsr*<0.1) in iPSC-CMs in hypoxia, we detect nearly identical enriched gene sets as for the DEseq2 results above. Among the genes that drive the Myogenesis enrichment, we found *COX6A2*, which is strongly upregulated in TAC lines (LFC=3.83) and is highly significant also in the DEseq2 analysis (adjusted *p* = 1.11E-07). It was recently shown that *COX6A2* deficiency in iPSC-derived CMs leads to impaired energy metabolism, abnormal calcium transients, and mitochondrial dysfunction, contributing to myocardial remodeling^41^. Interestingly, a set of *COX6A2* eQTLs in heart tissue from the GTEx project v. 8 (*p* = 4.6E-05) show substantial allele frequency differences between Tibetan and Sherpa populations and low altitude East Asians (Fig. 4), with the allele increasing expression being more common in Tibetans compared to CHB (i.e. 42% vs. 21%). This suggests that differences in frequency of cis-regulatory variants account for the higher *COX6A2* expression in TAC compared to CHB lines. Moreover, it points to protective cardiac traits that may be advantageous in the challenging hypoxic environment of high altitude.

**Fig 4.**
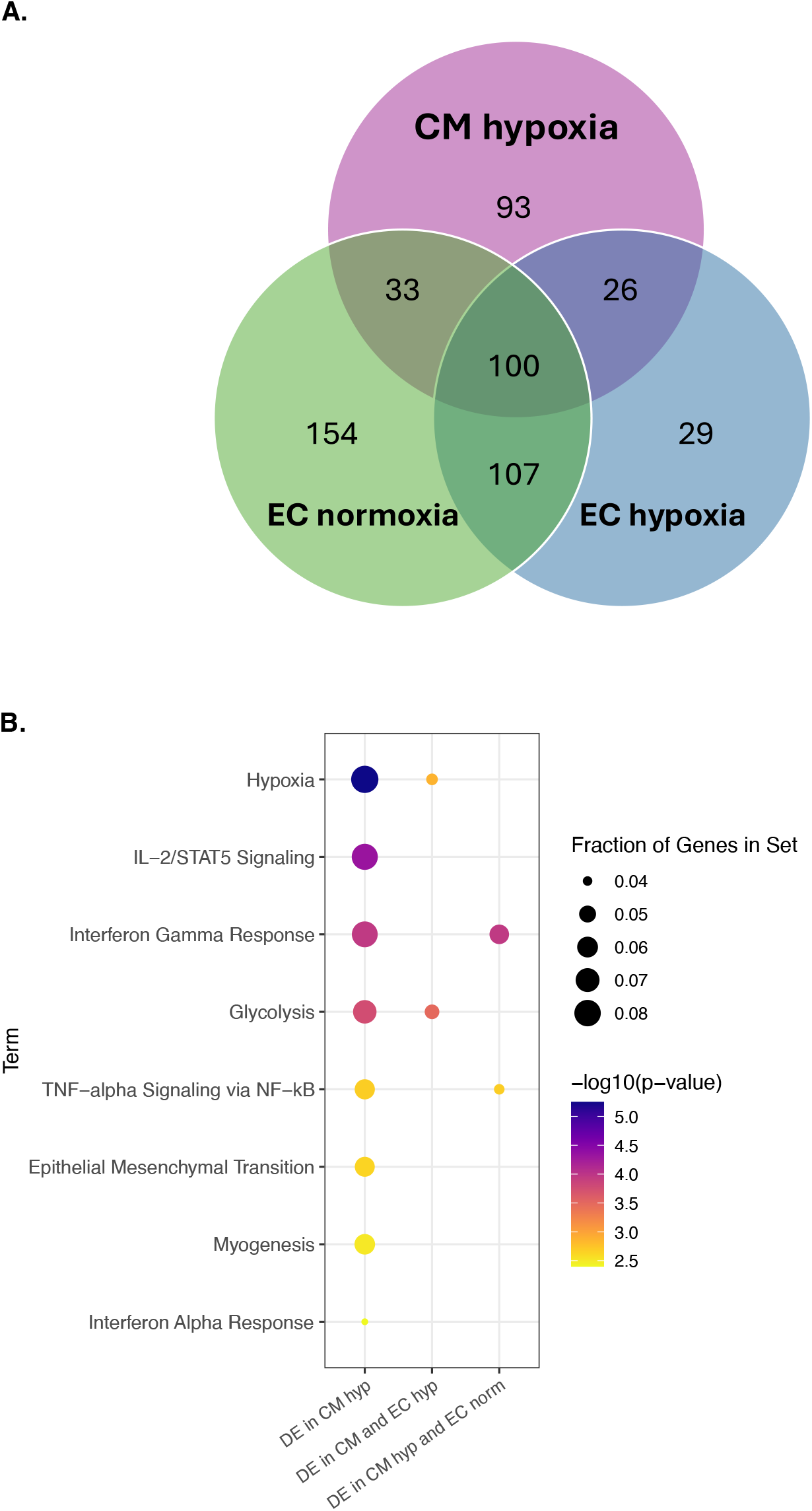
A) Venn diagram of the number of DE genes overlapping across iPSC-CMs in hypoxia, iPSC-ECs in hypoxia and iPSC-ECs in normoxia. The significance threshold for DE genes in each condition was *lfsr*<0.1. B) Gene set enrichment analysis of DE genes across populations in iPSC-CMs. This dot plot depicts the EnrichR results for all DE genes detected in the iPSC-CMs (*lfsr*<0.1) and corresponding results for the DE genes shared between iPSC-CMs (*lfsr*<0.1) and iPSC-ECs in normoxia and hypoxia (*lfsr*<0.1), respectively. Enriched terms are shown on the y axis. The gene set used in analysis is shown on the x axis.

Interestingly, when we focus on DE genes (*lsfr* <0.1) in both iPSC-CMs and iPSC-ECs in hypoxia, we find that only the Hypoxia and Glycolysis terms are significantly enriched (adj p<0.05), an enrichment that is likely driven by shared DE genes in the two terms (*GYS1, HSPA5, FAM162A, ENO2, IER3*, and *TGM2*). This is consistent with the profound impact that physiological hypoxia has on cellular energy production in many tissues, including the myocardium^42^. Interestingly, our previous work on iPSC-derived ECs had identified a subset of genes with an enhanced response to hypoxia in TAC compared to CHB cell lines. An example is the *ALDOC* gene which codes for an essential enzyme in the glycolysis pathway^43^ and is known to be upregulated in a hypoxic cellular environment^44^. We find that *ALDOC* is detected as upregulated also in TAC compared to CHB iPSC-derived CMs in both the DEseq2 analysis, which did not include the iPSC-ECs, and by mashr. Collectively, these findings suggest that the myocardium in Tibetans evolved specific metabolic adaptations to hypoxia; understanding these adaptations may have significant implications for the treatment of conditions such as coronary artery disease, myocardial infarction, and heart failure that decrease the amount of oxygen received by the heart.

In contrast, DE genes (*<u>lsfr</u>*<0.1) in both iPSC-CMs in hypoxia and iPSC-ECs in normoxia identify enriched pathways related to inflammation and immunity, i.e. Interferon Gamma Response and TNF-alpha Signaling via NF-kB, (adj p<0.05). Similar pathways were previously found to be enriched in iPSC-ECs in normoxia. No gene sets were significantly enriched in the DE genes (*lfsr*<0.1) in all 3 data sets, possibly due to the smaller number of genes and hence inadequate power.

In summary, our comparison of transcript levels in iPSC-derived CMs across TAC and CHB lines identified a number of interesting DE genes between populations and enriched gene sets that may underlie adaptive cardiac phenotypes in high altitude hypoxia. We previously reported DE genes and gene sets across populations in iPSC-ECs. However, we find that some genes are strongly DE only in the iPSC-CMs suggesting that Tibetans evolved adaptive traits in the vascular and cardiac systems. Our findings support the need for field studies comparing cardiac function across Tibetans and acclimatized lowlanders at high altitude.

## Materials and Methods

### Study Sample and Research Participants

The sample collection, establishment of LCLs, reprogramming to iPSCs and quality control tests are described in detail in Gray et al (2025)^21^. Briefly, Tibetan samples were obtained with informed consent (IRB16-1501) from Tibetans living in the Chicagoland area at the Tibetan Alliance of Chicago (TAC). The human subjects protocol was approved by the Institutional Review Board of the University of Chicago. All subjects were either born in Tibet or were the descendants of four Tibetan-born grandparents. Peripheral whole blood samples were collected by venipuncture, and LCLs were generated at the University of Chicago using a standard protocol (Coriell Institute) for establishment of LCL cultures from peripheral blood mononuclear cells. All Han Chinese samples used in the study were obtained from the Coriell Cell Repository in the form of LCLs. The samples were sex-balanced with equal numbers of male and female participants from each population group. The TAC samples were genotyped and imputed as described in Gray et al 2025.

Reprogramming of LCLs to iPSCs was performed in accordance with Burrows et al^45^ in batches, balanced by population and sex,. Standard quality control tests of pluripotency and stability were performed, including expression of endogenous and exogenous pluripotency factors, embryoid body assays to assess functional pluripotency, G-banded karyotyping using standard protocols^46^. No significant inter-population differences in gene expression in the iPSC lines was detected. All QC results are described in detail in Gray et al 2025^21^.

### Differentiation of iPSC-derived cardiomyocytes

Cardiomyocytes were differentiated as described in Knowles et al (2018)^26^ with a few modifications. Briefly, on day 0 feeder-free iPSCs were switched from essential 8 media into Heart Medium [RPMI 1640 without glutamine (Fisher, 21870-076)] supplemented with B27 -insulin (Invitrogen, A189560), glutaMAX (Life Technologies, 35050-061), and penicillin-streptomycin] supplemented with matrigel (0.1mg/ml) and 6μM of GSK3-inhibitor, CHIR990221 (Tocris, 4953). This was switched to plain heart media on day 1, then switched to Heart Media supplemented with 2μM Wnt-C59 (Tocris, 5148) on day 3 for 48 hours. Heart Media was refreshed on days 5, 7, 10, and 12. Wells were checked visually for contracting cells. Beating was typically observed between days 7 and 10. On days 14 – 20, wells with beating cells were purified using metabolic selection described in^47^. Cells were grown in Lactate Media [glucose-free RPMI 1640 (Gibco, 11879), with 500 ug/ml human serum albumin (Sigma-Aldrich, A0237), 213 ug/ml L-ascorbic acid 2-phosphate (Santa Cruz Biotechnology, sc228390), and 5% 1M lactate in HEPES (Sigma-Aldrich. L7022)]. Lactate is a metabolic substrate uniquely used by cardiomyocytes, particularly fetal cardiomyocytes^48^. By growing the cells in Lactate Media, refreshed every 48 hours, cells incapable of metabolizing lactate die and detach, while cardiomyocytes are enriched and purified. On Day 20, surviving cardiomyocytes were replated into Cardiomyocyte Maintenance Media (described in Burridge et al 2016^49^) which contains galactose (G5388, Sigma-Aldrich) rather than glucose as its primary energy source. The use of galactose forces the differentiated CMs to switch to aerobic respiration rather than anaerobic glycolysis. This shift mirrors the shift in metabolic program from fetal to adult heart cells and is intended to further mature the iPSC-derived cardiomyocytes. After a week in maintenance media, the cells were placed back in glucose-containing heart media and placed in hypoxia (1% O_2_, 5% CO_2_ – Coy Labs Hypoxia Glove Box) for 48hrs.

### Harvest and single cell sequencing

Following 48 hours of hypoxia exposure, cardiomyocytes were harvested as described in the 10x Genomics® Single Cell Protocols: Cell Preparation Guide (CG00053, Rev C). Cells were washed in DPBS and gently detached using Accutase, the accutase/cell mixtures were immediately placed on ice prior to removal from hypoxia chamber to prevent reperfusion injury and alterations to the transcriptional program. Cells were spun down at 4°C degrees and washed twice with cold 0.04% BSA in DPBS then filtered through 40uM filters to achieve single cell suspension. Cells were counted and assessed for viability then pooled at ∼250k-500k live cells per line. The pool was then washed a final time and resuspended in 1mL cold 0.04% BSA in DPBS. The pool was then transported on ice to the University of Chicago Genomics Core for quality assessment and library preparation using the 10x Chromium. Successful libraries were sequenced on the Illumina Novaseq (PE 100bp) (Table S8).

### Data Analysis

The cell ranger count pipeline of the 10X Genomics Cell Ranger 3.1.0 software package^50^ was used to align reads to the transcriptome (refdata-cellranger-hg19-3.0.0) and to generate matrices of UMI counts for each feature. To cluster the multiplexed scRNA-seq data by individual genotype and to identify potential doublets, we used the souporcell pipeline^51^ (build-date: Thursday_3_September_2020_2:51:20_UTC) which performs clustering without using reference genotypes. To map souporcell clusters to individual IDs, the souporcell cluster genotypes were compared to the genotypes of all 20 individuals at all overlapping positions with non-missing genotypes. For each souporcell cluster, a single individual from the demultiplexed batch was always found to have a genotype match rate of >70%, while the match rate was <50% for all other 19 individuals. The individual with the >70% match rate was identified with the cluster. Following cell clustering, the workflow of the scRICA 0.0.0.9000 R package^52^ was used to deconvolute doublets using the algorithm of DoubletDecon^53^.The output of scRICA was then imported into Seurat 4.1.0^54^ where the data were further filtered using the parameters of the standard pre-processing workflow (cells were retained if 1000 < gene count < 5000 genes and if MT count < 10%) and then normalized. The R package Harmony 0.1.0^55^ was used to integrate samples and correct for batch-specific conditions. Cell cycle scoring and regression was performed using the expression of G2/M and S phase markers following the Seurat workflow and using the list of cell cycle markers from Tirosh et al. 2015^56^. Seurat was then used to perform cell clustering using the Harmony embeddings. Cell cluster identification was based on the expression levels of 18 markers known to be associated with the 7 different cell types: CM/CM-mature, CM-precursor, atrial, ventricular, cardiac fibroblast (CF), mouse embryonic stem cells (MES) and iPSC^27,57^. Gene set enrichment analysis was also performed on the top 400 markers identified by the Seurat FindAllMarkers() function using the DAVID Functional Annotation Tool^58^ with the Human Protein Atlas Normal Tissue Celltype annotation database to provide additional validation of cell cluster identification. All R packages for single cell analyses were run in R 4.1.0.

Differential gene expression analysis between CHB and TAC lines was performed in a pseudo bulk manner by first aggregating read counts of those clusters identified as CM in the single cell analysis for each sample. Genes were then tested for DE using the R package DESeq2 v1.34.0^59^ with the design model ∼population+batch and the Wald test for significance. The final set of genes analyzed were those protein coding genes with at least 10 reads across all samples and those that passed the average expression threshold set by the independent Filtering function of DESeq2. P-values were corrected for multiple tests using the Benjamini-Hochberg method^60^.

All enrichment analyses were performed using EnrichR^37–39^ with a specified background. Briefly, EnrichR performs standard enrichment analysis and outputs a p-value using Fisher’s exact test, and an adjusted *p*-value using Benjamini-Hochberg correction for multiple testing. Results were reported as significant if the adjusted *p*-value was < 0.05. Hallmark MSigDB^61^ gene sets were included in analysis. For each independent analysis the background genes included all expressed genes in a given RNA-seq experiment.

## Data and Resource Availability

Further information and requests for resources and reagents should be directed to the corresponding author, Anna Di Rienzo. Cell lines generated in this study are available upon request. There are restrictions to the availability of the following cell lines due to conditions of study consent selected by research subjects: TAC086, TAC252, TAC540, TAC841. Genomic for the TAC samples and scRNAseq data for the TAC and the CHB samples are available through dbGAP (phs003758).

## Acknowledgements

We are grateful to all the members of the Tibetan Alliance of Chicago and, in particular, to all the donors of blood samples that made this research possible. We gratefully acknowledge: Yoav Gilad, Jonathan Burnett, Mengjie Chen, Michelle Ward, and Kenneth Barr for advice on experimental or analytical techniques. All single-cell RNA sequencing was performed at the University of Chicago Functional Genomics core (RRID:SCR_019196), which is supported by the Cancer Center Grant (P30 CA014599). This work was supported by a NIH grant to AD (HL119577); OAG was supported by NIH Training Grants (GM007197 and HL07605) and NIH F31 (HL142146).

## Author Contributions

AD, OAG, and DBW conceived of and performed the studies and analyses. Differentiation into cardiomyocytes and single-cell RNA sequencing were performed by OAG. All computational analyses were performed by DBW with support from OAG and AD. OAG, DBW and AD wrote the manuscript.

